# Dynamic land-plant carbon sources in marine sediments inferred from ancient DNA

**DOI:** 10.1101/2024.04.02.587465

**Authors:** Ulrike Herzschuh, Josefine Friederike Weiß, Kathleen Stoof-Leichsenring, Lars Harms, Dirk Nürnberg, Juliane Müller

## Abstract

Terrigenous organic matter in marine sediments is considered a significant long-term carbon sink, yet our knowledge regarding its source taxa is severely limited. Here, we leverage land-plant ancient DNA from six globally distributed marine sediment cores covering the Last Glacial-Holocene transition as a proxy for the share, accumulation rate, preservation, and composition of terrigenous organic matter. We show that the spatial and temporal plant composition as revealed by sedaDNA records reflects mainly the vegetation dynamics of nearby continents as revealed by comparison with pollen from land archives. However, we also find indications of a global north-to-south translocation of sedaDNA. The plant composition shows that upland vegetation is strongly underrepresented in the record compared to riverine and coastal sources. We also find that plant sedaDNA has a higher accumulation rate in samples from the Late Glacial, which is characterized by high runoff and mineral load. Thus plant DNA in marine sediments allows for new perspectives on the global linkages between the terrestrial and marine carbon cycle which would benefit from a more quantitative understanding of DNA preservation and dispersal. This represents the basis of how climate change and land-use change translate into carbon-sink dynamics and also informs about natural carbon-capture solutions.

## Introduction

Organic matter in marine sediments is recognized as a substantial long-term sink for atmospheric carbon dioxide (1). It is assumed that around a third of marine sediment carbon originates from different continental and coastal carbon pools, such as the biosphere and the soil (2). Land plants (embryophytes) including seed plants, ferns, and mosses account for the majority of the biomass on land (80–90 %) (3). However, we know little about which plant taxa form the terrestrial portion of the marine carbon sink and what the major source ecosystems and translocation pathways are. This restricts predictions on how carbon transport from source to sink is affected by ongoing climate and land-use change.

To link the terrestrial carbon sources with marine carbon sinks and their variations through time, we need a proxy that can depict the amount, degradation, and composition of land plant organic matter in marine sediments with high taxonomic resolution. Hitherto, even the separation of marine (4) and terrigenous (5) carbon has been a challenge for state-of-the-art biogeochemical and isotopic methods and only very rough taxonomic information has been available. For example, lignin-biomarker patterns have been used as proxies for the relative share of gymnosperms (6) and angiosperms (7); and fatty acid compound-specific isotope signals indicate the relative share and productivity of aquatic plants (8), but cannot differentiate between coastal and freshwater taxa. Furthermore, mixing models are applied to compound-specific isotope data to discern the relative contribution of C3 and C4 plants (9). In addition, pollen signals serve as indicators of the presence of terrestrial material in marine sediments and provide information on land changes even though they are restricted to seed plants and the biases of different pollen productivity and dispersal characteristics among the taxa need to be taken into account. Furthermore, pollen taphonomy in marine sediments differs strongly from the taphonomy of the majority of the vegetative plant biomass. Pollen is, at least partly, transported via air and is not preserved due to adsorption to mineral particles (10). Hence, it does reflect the linkages between the land and ocean carbon cycle only indirectly.

The analysis of ancient DNA (aDNA) found in sediments has revolutionized our ability to trace past changes in land ecosystems (11) and, recently, in marine ecosystems (12). In particular, sedimentary metagenomics – analyses of the total environmental DNA contained in sediments (13) – is a promising technique (14). In contrast to metabarcoding which relies on target-specific DNA amplification using PCR, non-target and PCR-free metagenomics has the advantage of being able to identify all major kingdoms quantitatively in a single approach (11). However, studies focusing on the terrigenous component in marine sediments are lacking. Even studies from land archives almost exclusively address ecological questions, while tracing the source of sedimentary organic carbon has rarely been the focus of aDNA studies (e.g., for permafrost) (15). Furthermore, it is believed that DNA preservation in the environment broadly resembles that of other organic compounds, where molecules are, to some extent, protected from degradation upon adsorption to minerals (16;17) i.e. not within cells of plant debris.

Here we present metagenomic analyses of six marine sediment records originating from the North Pacific (off-Kamchatka and the Bering Sea), the North Atlantic (Fram Strait, off-Svalbard), the tropical Atlantic (off-Tobago), off-Australia, and the Bransfield Strait (off-Antarctic Peninsula): all of them covering the last glacial–interglacial transition. Our analyses target the last Late Pleistocene-Holocene transition due to the known major changes in the terrestrial and marine carbon cycling in response to climate signals. We aim to identify the share, content, preservation, and taxonomic composition plant DNA. We seek to assess the spatio-temporal patterns of plant matter source taxa in marine sediment cores and compare them to land and marine proxy data. Our ambition is to trace the land-to-ocean carbon transport and marine carbon translocation as well as the terrestrial source pool and source ecosystems of plant matter in marine sediments. Finally, we discuss the potential of plant DNA in marine sediments as a proxy to link the terrestrial plant source with the marine carbon sink.

## Results

### Land-plant matter content and preservation

Metagenomic analyses of marine sediments yielded a substantial amount of plant DNA reads (Embryophyta) in all six globally distributed marine sediment cores (Fig. 1, 2). Shotgun sequencing and subsequent read filtering and taxonomic classification of 133 samples resulted in a total of 1,403,151 reads (DNA fragments) classified as Embryophyta, assigned to 242 taxa at the family level (Supplementary data S1, Supplementary data S2). Map damage-pattern analysis yielded the typical post-mortem C-to-T substitutions at the end of the reads, which increased with sample age (Supplementary figure 1).

**Figure 1.**
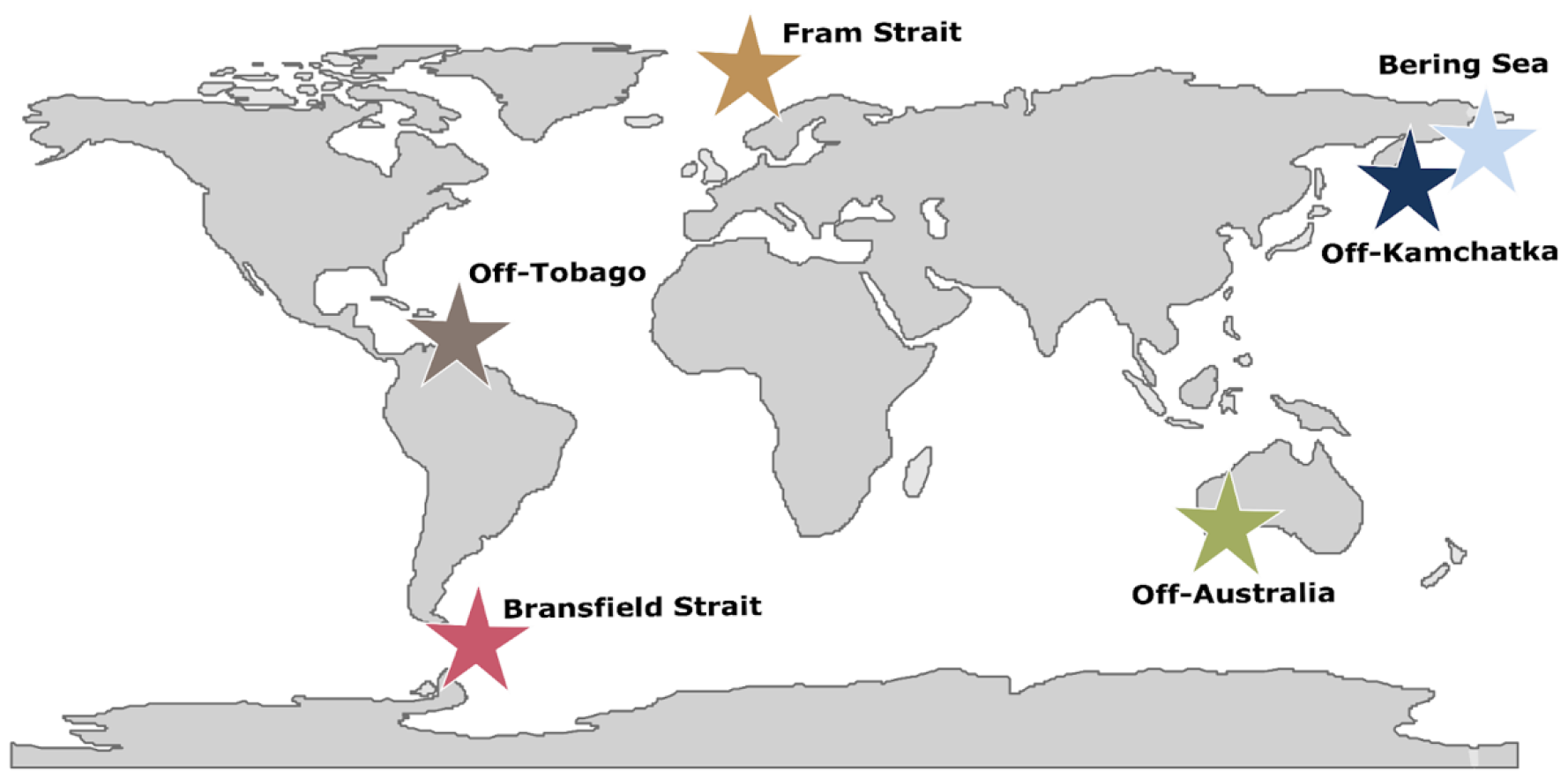
Map showing the location of the marine sediment cores: **Fram Strait** (MSM05/5-712-2, 78.915662 °N, 6.767167 °E), the **Bering Sea** (SO201-2-77KL, 56.3305°N, 170.6997°E), **off-Kamchatka** (SO201-2-12KL, 53.993°N, 162.37°E), **off-Tobago** (M78/1-235-1, 11.608°N, 60.964°W), **off-Australia** (MD03-2614G, 34.7288°S, 123.4283°E), and the **Bransfield Strait** (PS97/72-01, 62.006°S, 56.064°W).

**Figure 2.**
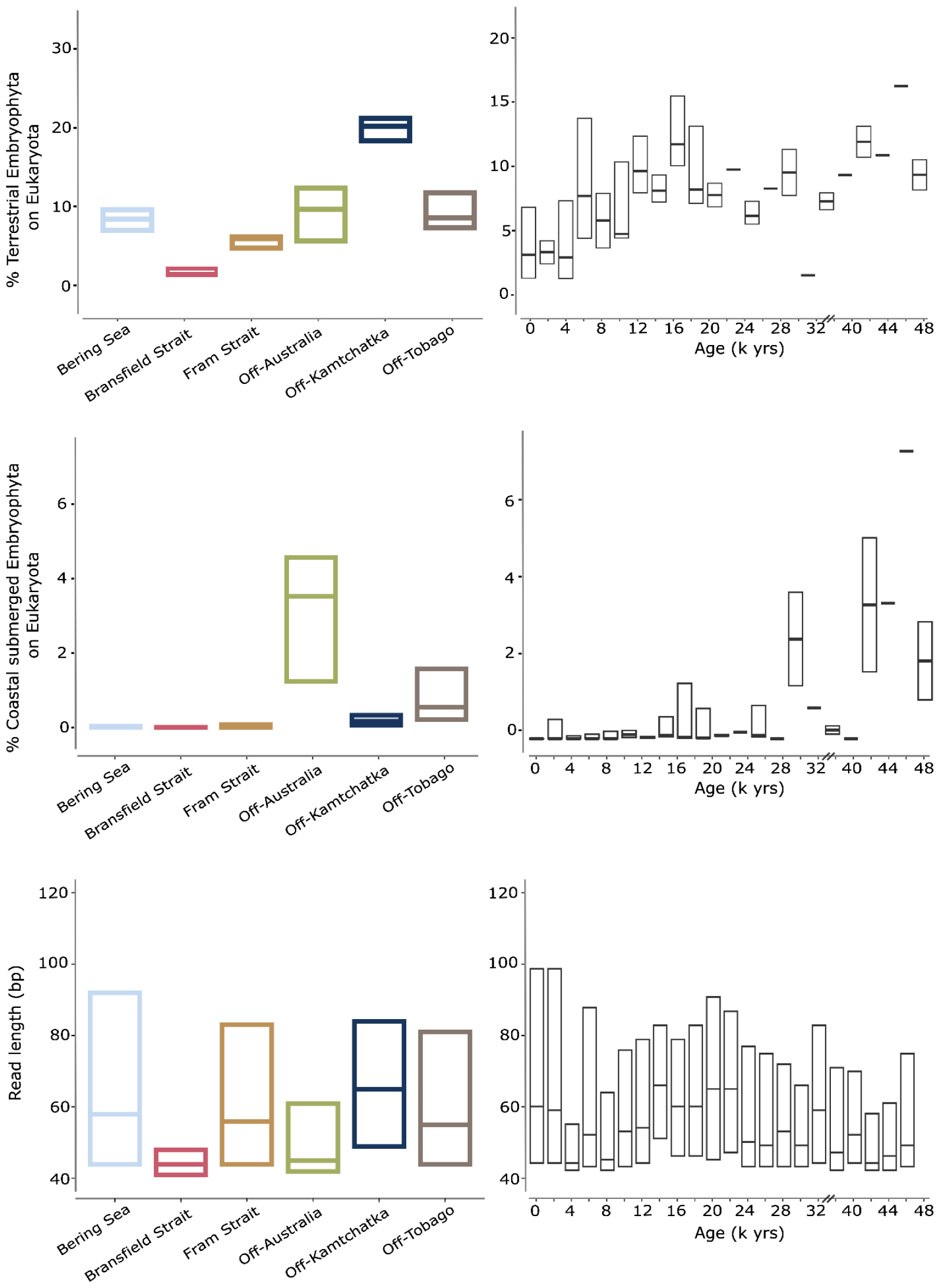
Family clade Embryophyta share (%) on total eukaryotic recovered sedimentary DNA and read length (bp) per core and per age (k yrs). Families are assigned to terrestrial vegetation (upper panels) or coastal submerged (lower panels). Embryophyta share (%) demonstrates the total Embryophyta reads (either terrestrial or coastal submerged) on the total Eukaryota reads.

The share of plant reads on all eukaryotic reads classified on family level ranges between 0.4 and 42,5% (median: 8.44, Supplementary figure 2) with most of them originating from terrestrial plants (Fig. 2). All cores show an increased terrestrial Embryophyta share during the last deglacial. The share of coastal submerged plants shows a different temporal pattern with higher values during the last glacial (Fig. 2). Only in the cores from Off-Australia and Off-Tobago coastal submerged Embryophyta form a major part (Fig. 2). This is the first time that the plant DNA share of marine samples has been reported. Our plant DNA results confirm the wide range of terrestrial organic matter contribution reported in biomarker studies (18). The share of land-plant DNA in samples from lakes (19) and permafrost soil (14) investigated using a similar methodology, is, as expected, higher.

The median plant DNA read length in the sediment cores ranges between 44 and 65 base pairs (bp). We interpret median DNA read length as a proxy for DNA fragmentation (20). Overall our results show a site-specific pattern with longest read lengths in the high-latitude Northern Hemisphere sites (Fig. 2). Median plant DNA read length in the individual cores does not decline with depth (Supplementary figure 2) suggesting that post-depositional signals do not dominate the preservation pattern which is in agreement with findings from terrestrial archives (21). Over all cores, median read length is higher during cold periods such as the LGM and Late Glacial and lower during warm periods such as the MIS 3 stadial and early Holocene (Fig. 2).

The plant DNA accumulation rate in the marine sediment core SO201-2-77KL (for which an accumulation rate was available) ranges between 0.0124 and 0.1 µg/cm^2^/yr (median: 0.08 µg/cm^2^/yr) which is of a similar order of magnitude to DNA accumulation rates in previous studies of surface marine sediment (22). The results from the sediment core show a clear temporal pattern with high values during the Holocene (Supplementary data S3), and at ∼14.6 and ∼11.5 k yr BP (Fig. 3), reflecting known phases of increased terrestrial productivity and terrestrial organic matter (OM) delivery by meltwater runoff (23; 24). Furthermore, the accumulation of land plant DNA shows the same peaks as the Lignin accumulation rate of the marine sediment core SO202-18-3/6 that is located close to SO201-2-77KL in the Bering Sea (7).

**Figure 3.**
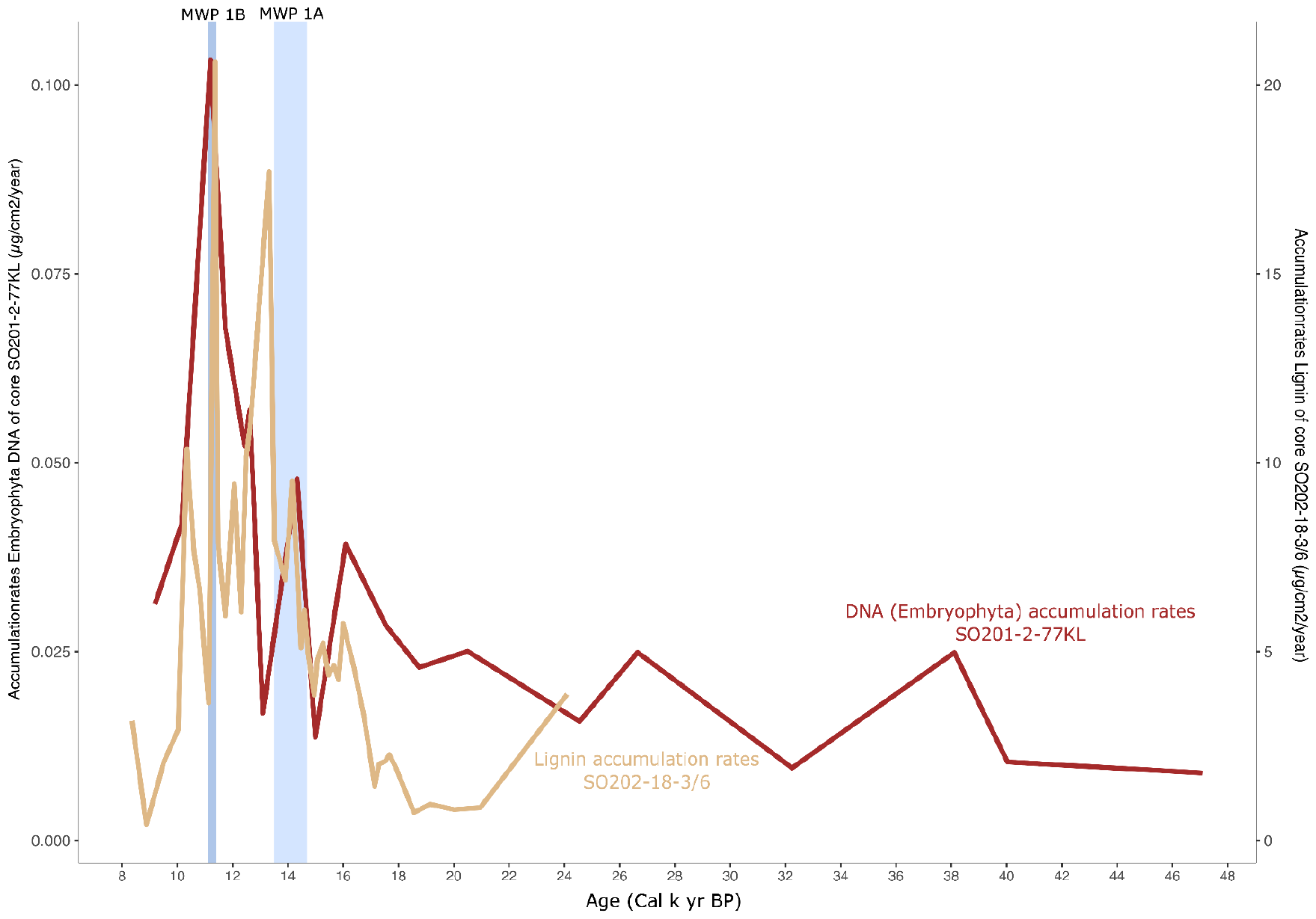
Comparison of Embryophyta DNA accumulation rate (µg/cm^2^/yr) for the marine sediment core SO201-2-77KL and the Lignin accumulation rate (µg/cm^2^/yr) of a proximal marine sediment core (SO202-18-3/6). Both cores are located in the Bering Sea. The lignin accumulation rate was published by Cao et al., 2023.

### Major compositional pattern of land-plant DNA

Principal component analysis (PCA) is used to show the similarity of the samples in terms of plant taxa composition. Figure 4 visualizes the direct grouping of samples from the same sediment core. In addition, the ordination shows that the plant DNA composition of the different ages of individual sediment cores are largely characterized by plant families known from the flora of neighboring continents (Fig. 4).The arctic and subarctic records (Fram Strait, off-Kamchatka, Bering Sea) cluster with typical arcto-boreal families including Saxifragaceae, Cyperaceae, Betulaceae, Rosaceae, Boraginaceae, and Salicaceae. The Bransfield Strait record (off-Antarctic Peninsula) is located close to and therefore mainly characterized by moss families, for example, Marchantiaceae and Polytrichaceae (Fig. 4, 5). Although Cyathodiaceae are not known from Antarctica, they may reflect liverwort abundance in general, as liverworts are poorly covered in the reference database. Other taxa not native to Antarctica characterize the Bransfield Strait plant spectrum; however, these are common in Patagonia (e.g., Cupressaceae) or have a characteristic distribution in the Southern Hemisphere (e.g., Restionaceae, Hymenophyllaceae). The samples of the record coming from off-Tobago show the accumulation of many tropical plant families including Myrtaceae, Melastomatacae, Rubiaceae, Moraceae, Chrysobalanaceae, and Lauraceae. The record is also unique in its relatively high share of Rhizophoraceae representing typical members of the tropical mangrove forest (Supplementary figure 3). The record from off-Australia is characterized by a high number of tropical seagrass families such as Cymodoceaceae and Posidoniaceae which are typical of southern Australia’s sandy coasts as is shown by their close proximity in Figure 4 (25). Families endemic (Maundiaceae) or common (Asparagaceae) to the Australian region also occur.

**Figure 4.**
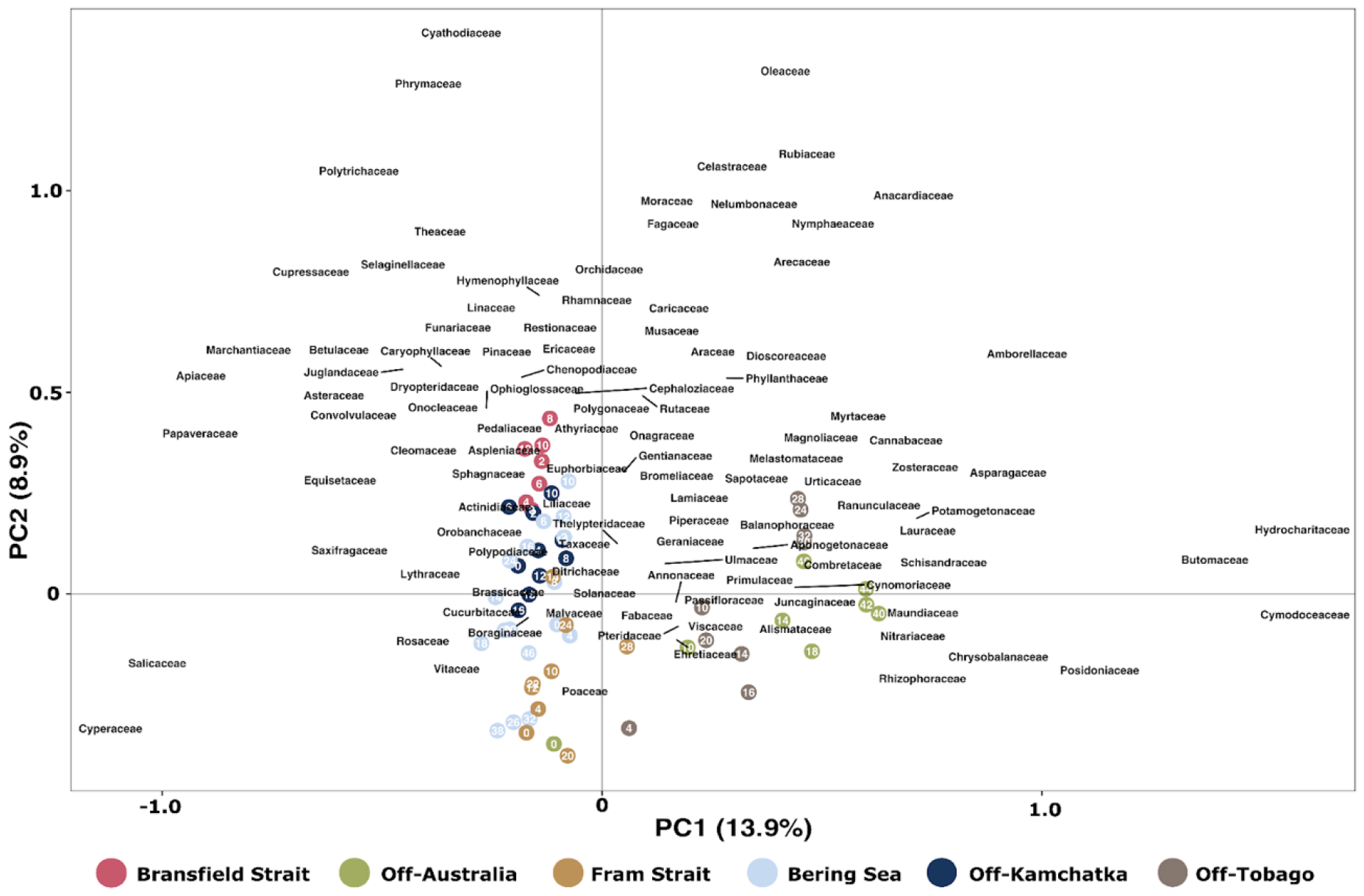
Principal component analysis of the plant DNA composition in the timeslices of the six marine cores. The scaling was set to 1 and only taxa with a minimum relative abundance of 0.5% and an occurrence in 3 time slices are shown. To avoid overlapping, not all plant families are displayed. The plot indicates that plant DNA composition is mainly site-specific (samples from each core cluster together) but also show temporal trends (e.g. the consistent temporal variation of the Bering Sea and Off-Kamchatka record along the axis 2). Generally taxa with particularly high or low axis values are more distinctive for the composition of single samples (i.e. those placed into the same direction) while taxa placed in the center are common among all samples.

**Figure 5.**
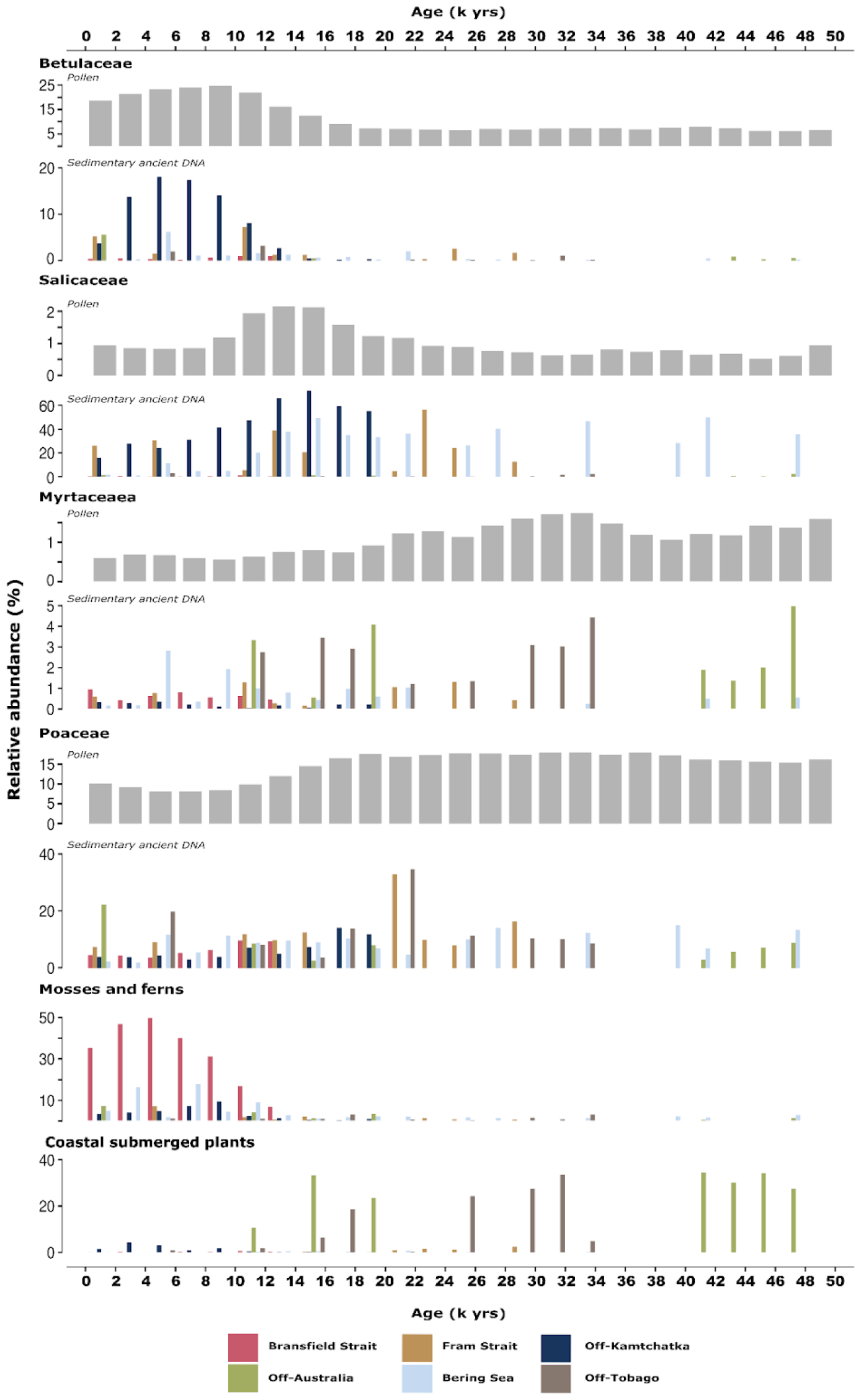
Stratigraphic plot of plant DNA composition (relative abundance of all reads ranked on family level (%)) in six marine cores over the last 50,000 years compared to pollen data synthesized from terrestrial archives (28). For Betulaceae, Salicaceae, Myrtaceae, and Poaceae the gray bar graphs show the global share of each taxon as % of all terrestrial pollen taxa (see methods). Moss and fern families were grouped together as well as the coastal submerged plants.

Typical crop plant families such as Solanaceae, Malvaceae, Bromeliaceae, Brassicaceae, and Vitaceae are placed in the center of the ordination plot (Fig. 4). Read length analyses indicate that reads assigned to these taxa are mostly short (Supplementary figure 4) and thus have a higher chance of being assigned to taxa overrepresented in the database like crop plants. Accordingly we assume that these assignments are false-positives due to a highly degraded unspecific DNA component shared by all samples. Such false classifications can result from short sequence length, which reduces the diagnostic sequence differences that are needed to gain an exact taxonomic classification, and by the overrepresentation of genomes in the reference databases, which increases the likelihood of read-reference hits.

## Discussion

### Spatio-temporal pattern of plant matter source taxa in marine sediment cores compared to land and marine proxy data

Major shifts in the plant composition in the marine DNA record align with corresponding pollen taxa changes in the source area of the land plant DNA found in continental paleo archives (Fig. 5). Grass and herb taxa such as Poaceae, Cyperaceae, and Papaveraceae show higher values around the Last Glacial Maximum, approximately 25,000 to 19,000 years ago, in the northern high latitude records suggesting an intensified contribution of biomass from widespread Northern Hemisphere glacial steppes (26) to marine carbon burial. A heightened contribution of C3 plants, the preferred metabolic pathway of Poaceae that are adapted to dry cold climates, was discerned in high latitude marine sediments of the Last Glacial period (see Fig. 5 and Supplementary figure 10). Furthermore, the glacial samples contain abundant reads from seagrass families (>20 %) (Fig. 5), likely due to lower sea levels when shelf areas were more densely covered, but may also relate to a taphonomic signal because shallow areas were located closer to the coring sites (27).

The transition from the glacial to the deglacial phase introduces notable changes in the marine sedimentary DNA record. Increased terrigenous organic matter sedimentation is indicated by the plant DNA share and plant DNA accumulation rate in cores off-Kamchatka and the Bering Sea, aligning with biomarker studies which show a higher input of angiosperm woody taxa during the deglacial (7) (Fig. 3 and Supplementary figures 6, 7). Specifically, Salicaceae (*Salix, Populus*), the willow family, peaking at 14,000 years ago (Fig. 5), represents a dominant contributor to land-plant carbon in marine sediments in the northern high latitudes during this period, confirming the global Salicaceae pollen trend (28). It most likely reflects the riparian woodland expansion in response to enhanced water supply from glaciers melting during the Late Glacial, peaking at the temperature rise at the beginning of the Bølling/Allerød period (29). Likewise, the core in the Fram Strait shows a peak during the Glacial–Holocene transition in the Salicaceae and Poaceae families, which is consistent with studies that also observe high levels of terrestrial organic matter during this time in this marine region (Fig. 5) (30). In accordance with a regional glacial discharge event as the primary driver, we see increased terrestrial plant DNA input in the Bransfield Strait (Supplementary figure 10). The low latitude sites demonstrate substantial compositional turnover during the Late Glacial–Holocene transition as well, with Myrtaceae, Rhizophoraceae, and Chrysobalanaceae DNA showing elevated values during the Late Glacial and early Holocene, potentially indicating intensified run-off processes as well (Fig. 5, Supplementary figure 3) (31). This inference is substantiated by biomarker evidence indicative of mangrove extent (Supplementary figures 3, 7, 8). The sediment cores off Australia show a transition from coastal submerged plants to terrestrial regional plants associated with a rise in sea level from the Glacial to the Holocene (Supplementary Figure 10).

**Figure 6.**
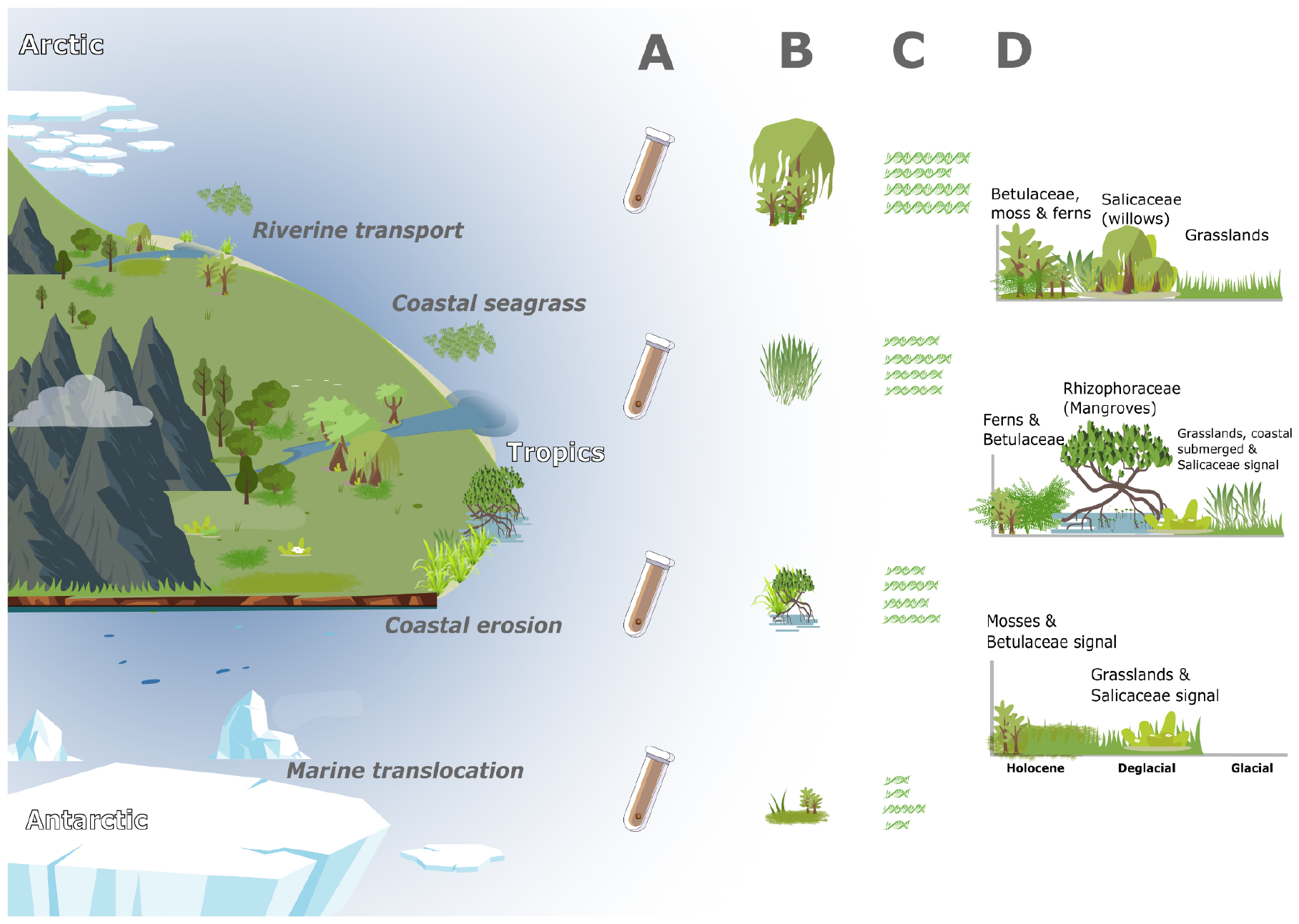
Transport of organic matter from land plants and insights into ancient DNA. This illustration shows the major transport pathways for organic matter from land plants into the marine ecosystem on the left. The different transport pathways can be seen in all geographic zones and at all latitudes. The right panel illustrates what this research has revealed by detecting ancient DNA from terrestrial plants in marine sediments and summarizes our main findings. **A** illustrates that the sediment samples were taken from different geographical settings. **B** shows that the different geographical settings show a difference in the amount of ancient land-plant DNA in the marine sediments based on the Embryophyta to Eukaryota ratio (%). **C** demonstrates that the read length of the recovered ancient DNA shows a spatial pattern based on length of transport of Embryophyta DNA. **D** illustrates the compositional changes in land-plant organic matter sources from the Northern Hemisphere over the Tropics to the Southern Ocean. The pictograms used in this illustration were designed by Freepik.

The Holocene period brings further changes to the marine sedimentary plant DNA record. Betulaceae emerges as a strong contributor to the land-to-ocean plant DNA flux in the northern high latitude marine sites, reflecting the massive expansion of this family on the nearby continents in response to early Holocene warming (Fig. 5). There is also a significant presence of Betulaceae DNA in tropical and Southern Hemisphere records (up to 4%), which are remote from their typical distribution area (Fig. 5). Additionally, ferns and mosses show extended DNA contributions in all records since the beginning of the Holocene, except for the tropical record from off-Tobago (Fig. 5). The presence of ferns and mosses during the early Holocene in the plant DNA records from off-Kamchatka and the Bering Sea confirms land palynomorph data (32).

Our plant DNA data indicate a successive increase in moss share in the Bransfield Strait record from 12,000 years BP onwards (Fig. 5). This may point to an earlier than expected retreat of the Antarctic Peninsula Ice Sheet from the northern part of the peninsula (including the South Shetland Islands), which is estimated to have occurred 10,000 years ago (33). Generally, a moss share of up to 50% of plant DNA in the Bransfield record during the mid-Holocene warm period confirms increased terrestrial input as inferred from biomarker studies in the sub-antarctic (34).

Overall, our observations identify the source taxa that contribute to the land plan DNA record in marine sediments indicating the potential of the proxy to unveil the terrigenous carbon source in the marine sediments and their spatial and temporal variations. Moreover, this marine sedimentary proxy provides insights into the vegetation dynamics on land and in near-shore areas.

### Land-to-ocean carbon transport and marine carbon translocation

Spatial and temporal patterns in plant DNA share, read length, accumulation rate and composition suggest that the land-plant carbon amount, preservation, and source in marine sediments is mainly a function of organic matter supply via rivers and meltwater input from nearby continents. The sites off-Kamchatka and off-Tobago, for example, have the highest mean shares in land plants in accordance with their proximate locations to the Kamchatka River and Orinoco River estuaries, respectively (Fig. 2). In contrast, the coring sites from Fram Strait (off-Svalbard), Bering Sea, and Bransfield Strait (off-Antarctic Peninsula) have low shares of reads assigned to Embryophyta reflecting their remote location to terrestrial plant sources (Fig. 2). The riverine impacted locations also have longer median read length compared with remote sites from a similar latitude. Furthermore, ordination results portray a distinctive plant composition for each sediment core, characterized by taxa known from the regional flora (Fig. 4).

At a global scale, the spatial differences in the taxa composition are stronger than the temporal changes within the records. This interpretation is supported by nearby records from the Bering Sea showing similar compositional trends. These results suppose a spatial consistency between the nearby continental plant DNA source and the marine sink. Furthermore, we find that the deglacial, which is known as a period of high continental run-off and mineral load (31), is characterized by a higher and less degraded plant DNA share, an increased land-plant DNA accumulation rate, and an enhanced contribution of riverine taxa compared with Holocene samples (except for Bransfield Strait) (Fig. 2; Fig. 3; Fig. 5 and Supplementary figure 2). Thus, our plant DNA signals propose that the transport of terrigenous material via rivers toward the shelf regions translates continental source dynamics – which are sensitive to climate change – into marine carbon sink dynamics (35).

The observed plant DNA degradation (based on read length distribution) and composition suggest a north-to-south carbon translocation at a global scale via large-scale ocean currents which has rarely been investigated using data evidence. We find that the plant DNA of the Arctic Fram Strait record is much less degraded (median: 55 bp) compared with the Antarctic Bransfield Strait plant DNA (median: 44 bp) (Fig. 2, Supplementary figure 2) despite their comparable geographic settings. Furthermore, we observe clear signals of Betulaceae in the tropical and Southern Hemisphere cores despite their remote location from the natural distribution area of Betulaceae on the proximal continents (Fig. 5). The signals in most of these sites increase during the Holocene, as seen in the northern high latitude records and the pollen signal. This may represent the first plant compositional signal of broad-scale plant carbon translocation. Our data confirm modeling approaches which support a strong link between organic carbon translocation and thermohaline circulation, potentially indicating a strong Northern Hemisphere contribution to the Southern Hemisphere carbon sink (36).

### Terrestrial source pool and source ecosystems of plant matter in marine sediments

The broad agreement between the ancient plant DNA in marine sediments and land archive derived pollen-based trends on a millennial time-scale indicates that the plant DNA signal is mainly derived from an immediate, or at least not severely pre-aged carbon pool. This means that plant DNA can serve as a proxy for the burial of recent plant matter and such for carbon sink on human-relevant timescales. In contrast, compound-specific radiocarbon studies indicate that common biomarkers for terrigenous matter, such as leaf waxes and lignin, originate, at least partly, from a strongly pre-aged organic carbon pool, for example, from permafrost soil (7), though biomarker-specific turnover times have been recognized (38). Hence, the lateral transport of old soils or sediments (37) may be less reflected in the plant DNA signal but this requires further research.

While the temporal patterns show many similarities between the marine sedimentary aDNA and pollen records from terrestrial archives, the relative taxa shares show marked differences (Fig. 5). This confirms the different taphonomies of sedimentary pollen (via air) and sedimentary DNA (via erosion of plant organic matter) (39). Compared with upland ecosystems, the lowlands, in particular the riverine and coastal woodlands, are likely a major carbon source due to their direct connectivity with the land-ocean pathways (40). This is exemplified by the overrepresentation of Salicaceae and Rhizophoraceae in the marine sedimentary record compared to their role in the terrestrial vegetation (Fig. 5; Supplementary figure 3). Our data show a surprisingly low contribution of Pinaceae and Fagaceae to the record despite them forming an important plant carbon pool in the northern mid-to-high latitudes (41) and are comparatively well represented in the reference databases. This finding is independent evidence that transport efficiency (e.g. erosion of riverine vegetation) rather than on-land productivity determine the carbon sink of land-plant matter in marine sediments (2).

### Plant DNA in marine sediments as a new proxy to link the terrestrial source with the marine carbon sink

The obtained plant DNA proxy information from the six globally distributed records covering the Last Glacial–Holocene transition yielded spatially and temporally distinct signals which we interpret with respect to changes of the share, preservation, and composition of terrigenous plant DNA in marine sediments. We are confident that the obtained DNA signals hold useful information about the share, preservation, and composition of terrigenous organic matter (Fig. 6) because the obtained results match major expectations about changes in the land source signal gained from pollen studies on land archives. For further validation and assessment purposes, we conducted a comprehensive comparative analysis of our plant-aDNA derived signals and state-of-the-art terrestrial biomarkers and other proxies indicative of terrigenous material present in the same or nearby marine sediment cores which yielded corresponding trends (Supplementary figure 5, 6, 7, 8, 9, 10). However, state-of-the-art proxies can only roughly indicate the source material, for example, share of woody taxa and the angiosperm/gymnosperm ratio (5), while we provide information on the source taxa at family level representing a hitherto unparalleled taxonomic resolution. For example, we find temporally contrasting organic matter contributions from Salicaceae and Betulaceae. Because of this, we are able to trace a distinct contribution of taxa restricted to the Northern Hemisphere in Southern Hemisphere sediment cores. Also a clear distinction of the temporal trends of the major plant functional types (i.e. moss, ferns, herbs, and woody taxa) within terrestrial material has not yet been documented. We are the first to report the contribution of coastal submerged plants to the marine sedimentary organic matter, which we find to be a substantial, albeit temporally and spatially highly variable, portion (Fig. 6).

With the ongoing completion and curation of the reference databases and further development of bioinformatic pipelines (11), the quality of the results obtained in future studies will be improved. Additionally, recent studies highlight that organic matter preservation is dependent on the strength of the organo-mineral bonds rather than selective preservation of different organic compounds (17). This implies that sedimentary ancient DNA signals can serve as a proxy not only for the amount, preservation, and taxonomic origin of sedimentary plant DNA but, to some extent, also of the total plant organic matter signals. Our spatiotemporal pattern of DNA read length would imply that organic matter preservation and as such the marine carbon sink for terrestrial material on a millennial time-scale is mainly influenced by pre-depositional (climate, mineral supply) rather than post-depositional processes (42). However, before plant DNA in marine sediments can be used as a reliable proxy for a substantial portion of terrestrial organic matter in marine sediments, it requires a better understanding of how and where DNA stabilization via organic-mineral particle formation occurs along the transition from fresh plant material to litter and soil. Also further studies are required to assess the DNA dispersal characteristics in comparison to those of other types of terrestrial organic matter (43).

To conclude, our results indicate that plant DNA in marine sediments can reveal the connection between terrestrial plant sources and marine sedimentary sinks for terrestrial carbon. Based on this knowledge this proxy could ultimately unveil the major carbon pools, disclosing transport pathways from land to ocean, and shedding light on terrestrial carbon translocation within the ocean itself (Fig. 6). This allows for new perspectives on the global linkages between the terrestrial and marine carbon cycle. Future studies on high-resolution archives can also inform how recent climate and land-use change has impacted the marine sink. The proxy also has the potential to indicate risks as well as to assess the potential of natural carbon capture solutions (4).

## Materials and Methods

### Sediment Cores

Six marine sediment cores were analyzed for this study. During cruise SO201 Leg 2 in 2009, core SO201-2-77KL (Shirshov Ridge, 56.3305°N, 170.6997°E) and core SO201-2-12KL (off Kamchatka, 53.993°N, 162.37°E) were recovered from the Bering Sea, detailed descriptions and the age depth model can be found in references 44 and 45. The shotgun DNA dataset of the core SO201-2-12KL is published by reference 12. For SO201-2-77KL the shotgun DNA dataset is published by reference 46. Core MSM05/5-712-2 comes from the Fram Strait (78.915662 °N, 6.767167 °E) (47). The age depth model was published in reference 47. A core from the Tobago Basin M78/1-235-1 (11.608°N, 60.964°W) represents the subtropical region. Main information on this core including the age depth model can be found in reference 48. A core near the southern Australian coast named MD03-2614G (34.7288°S, 123.4283°E), represents the Southern Hemisphere (49), as does the last core, designated PS97/72-01, which was recovered from the Bransfield Strait at the northern tip of the Antarctic Peninsula (62.006°S, 56.064°W) (50). For the core MD03-2614G the age depth model was published in reference 49 and for the core PS97/72-01 it was published in reference 50. Sediment cores were selected from sites at high and low latitudes, in the northern and southern hemispheres, distal or proximal to estuaries, considering the availability of records with reliable age-depth models and accessibility. Distal locations in particular show the potential of sedaDNA to track the terrestrial signal from the coast to more offshore locations.

### Extraction of sedaDNA

PS97/72-01 and MSM05/5-712-2 were subsampled in a clean climate chamber at the GFZ Potsdam, Germany. Samples of the cores SO201-2-KL77, SO201-2-KL12, M78/1-235-1 and MD03-2614G were taken in a subsampling room at the GEOMAR, Kiel, Germany. Avoiding contamination by modern DNA, all subsequent working steps were performed in a dedicated ancient DNA laboratory at AWI Potsdam and control samples were used to monitor lab contaminations. In total, DNA from ∼126 sediment samples was extracted using the DNeasy PowerMax Soil Kit (Qiagen) and the DNA was purified and concentrated using the GeneJET PCR Purification Kit (Thermo Fisher Scientific). Finally, DNA was quantified and diluted to 3 ng/µL.

### Shotgun metagenomics

DNA samples were prepared for whole genome sequencing using a specified single-stranded DNA library protocol (51) using an input of 30 ng per sample. For sample demultiplexing DNA libraries were indexed, pooled equimolarily, and shotgun sequenced using a NextSeq 2000 system (Illumina) with a customized sequencing primer CL72 in paired-end mode (2x100 bp), except for DNA samples from core SO201-2-KL12 (12) that were sequenced on a HiSeq 2500 system (Illumina).

### Bioinformatic processing

After sequencing, the raw sequencing reads were processed in a bioinformatic pipeline, which includes quality checking with FastQC (version 0.11.9) and adapter trimming and merging using Fastp (0.20.1). The taxonomic classification of passed sequencing reads was conducted with *Kraken2* (52) at a confidence threshold of 0.5 against the non-redundant nucleotide database, which was downloaded in April 2021 and converted into a *Kraken* database with a k-mer size of 35 bp and a minimizer size of 31 bp.

The *Kraken2* report files were then processed using a Python script to add the full taxonomic lineage information. Additionally, the read length was analyzed using the Fastp and Kraken2 files. The read length was then analyzed in R (R Core Team, 2022).

### Post-mortem damage pattern analysis

Post-mortem damage patterns are DNA signatures indicating the authenticity of ancient DNA molecules. Typically, an increase in the nucleotide frequency of C (Cytosine) to T (Thymine) at the end of the sequencing reads compared with a genomic reference is evident. For the analysis of post-mortem damage patterns, we extracted *Salix* sp. (taxID 40685) reads from the sequencing data of the core SO201-2-12KL, which were grouped into three datasets (ds01: 1081–5600 years BP; ds02: 6200–12,600 years BP; ds03: 13,600–19,900 years BP).

Thereafter, initial sequencing files were consolidated using the cat command, followed by filtering, merging, and taxonomic classification using *Kraken2* against the nt database with a lower confidence threshold of 0.2 compared to the previous classification (0.5) to increase the number of Salicaceae reads as initial input for the analyses. Then, *Salix* reads were extracted and mapped against the reference genome (*Salix brachista*, Accession number: GCA_009078335.1) in MapDamage (v. 2.0.8) (53) using the options --rescale (to adjust quality scores for potential misincorporations resulting from ancient DNA damage) and --single-stranded (to accommodate a single-stranded protocol). C to T frequency changes at the ends of the DNA reads are plotted in fragment misincorporation plots together with the fragment length distribution (Supplementary figure 1).

### Statistics, calculations and plotting

Further statistical analyses and plotting of data was performed in RStudio (R version 4.2.2; R Core Team 2022). All graphics were exported as vector graphics (e.g., PDF) to enhance them afterwards in the graphic design program Inkscape (Inkscape 1.1.2 (b8e25be8, 2022-02-05)). Pictograms for illustrations were designed by Freepik (www.freepik.com). For better comparison, all datasets were grouped in 2000-year time slices.

The initial step of handling the ancient DNA datasets involved reading and structuring data from separate files using the R packages *data*.*table* (54), *stringr* (55), and *tidyverse* (56). The read dataset was imported from a combined *Kraken* report file. Subsequently, taxonomic lineage information was merged with the initial dataset and the metadata were included as well. The taxonomic focus narrowed to the Eukaryota superkingdom, and further steps isolated data on the Streptophyta phylum, which after exclusion of specific non-embryophyta algae families (Chlorokybophyceae, Klebsormidiophyceae, Mesostigmatophyceae, Charophyceae, Coleochaetophyceae) resulted in a dataset with only Embryophyta. Within Embryophyta, several subgroups were examined, specifically identifying embryophytes associated within and outside of coastal areas or moss and ferns. Total clades were calculated for each subgroup in relation to different age ranges. The analysis of the proportion of Embryophyta and the subgroups of the total eukaryotic reads involved grouping the data by age, and then calculating various ratios (in %) within each age group. The calculated values were added as new columns to the dataset.

The read length analysis was conducted in RStudio using the taxonomic identification derived from the *Kraken2* pipeline (52) for each read per sample for each core using the R package *stringr* (55) and plotted using *gridExtra* (57). Each file was iteratively processed and partial file names were matched to entries within a results table. Additionally, age values were linked with corresponding file names. The accumulated data were methodically incorporated into the evolving results data table. We assessed the normality of the data distribution and tested for variance homogeneity prior to conducting the analysis of variance. While the data demonstrated a normal distribution, the variance homogeneity assumption was violated. Differences in read length per core were then evaluated using Welch’s test and the Games-Howell post-hoc test using the function *anovaOneW* in the R package *jmv* (Supplementary figure 11) (58).

The sediment accumulation rate was calculated using the dry and wet bulk densities and the linear sedimentation rate according to Riethdorf et al. (2013). Generally the total DNA accumulation rate was calculated and relative abundance of Embryophyta read counts was used to estimate the Embryophyta DNA accumulation rate. First, we calculated a *cumulative concentration factor C* and a *cumulative dilution factor D, which* sum up all concentration and dilution steps from DNA extraction until DNA sequencing. Second, the final weight of total DNA (*DNA*_*Final*_ *(g))* that has been sequenced was calculated by multiplying the final concentration *c*_*Final*_ *(pM)*, the molecular weight of a nucleotide *M*_*Nucleotide*_ *(325 pg/pM)* and the *average fragment length (bp)* of the DNA library used for DNA sequencing.

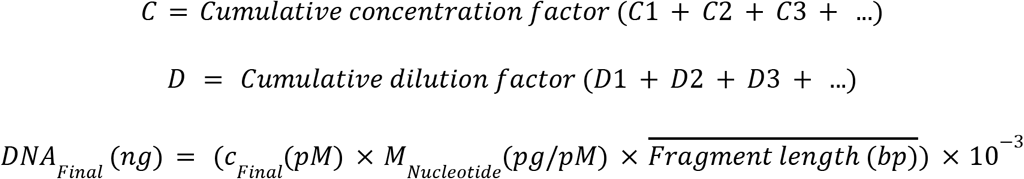

To obtain the initial total DNA weight per sample, we multiplied the *DNA*_*Final*_ *(g)* by the *cumulative dilution factor D* and divided by the *cumulative concentration factor C*.

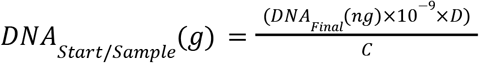

Finally we estimated the DNA accumulation rate for dried sediment samples. Therefore we used the known water content, calculated using the Wet Bulk Density (WBD) and the Dry Bulk Density (DBD), of the sediment samples and calculated dry weights for the samples used for the sedaDNA analysis. The weight of DNA per gram of sediment before processing the sample *DNA*_*Start*_ *(wt %)* is calculated by dividing the *DNA*_*Start / Sample*_ *(g)* by the dry weight of the sample *m*_*Sample*_ *(g)*.

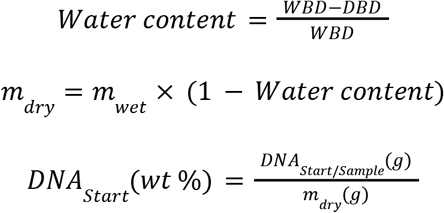

The sequencer used for our samples has a sequence coverage of 99%, so we know that the entire sequences are covered (Manual NextSeq 2000 system, Illumina). Therefore, the percentage of a particular taxa group, in our case Embryophyta, was calculated per sample and our *DNA*_*Start*_ *(wt %)* was multiplied with the percentage to get *Embryophyta DNA*_*Start*_ *(wt%)*.

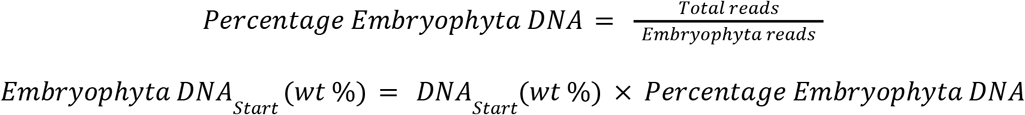

We were then able to calculate the accumulation rate of Embryophyta DNA using *AR*_*WDB*_ and the weight of Embryophyta DNA per gram of sediment, *Embryophyta DNA*_*Start*_ *(wt %)*.

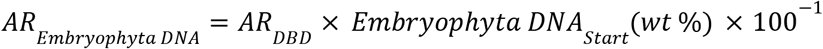

Characteristic families in the marine sediment cores were plotted against terrestrial pollen records from a global taxonomically harmonized pollen dataset (28). The relative abundance of the DNA record was calculated per core per time slice using the counts of Embryophyta that are at least classified to family level (Rank=F). Pollen percentages of samples were first aggregated per time-slice, then per region (Asia, Europe, North America, South America, Africa, and Indopacific) and then averaged globally. For plotting we used *ggplot2* (59), *tidyverse* (56) and *tidypaleo* (60).

We ran a principal component analysis using the R packages *analogue* (61), *stringr* (55), *ggrepel* (62), and *ggplot2* (59) for all cores together. Only taxa with an occurrence in at least 3 time slices and a minimum relative abundance of 3% were included.

## Acknowledgments

We thank Thomas Böhmer for his help in preparing the pollen dataset. Furthermore, we acknowledge Cathy Jenks for manuscript proofreading.

