## Supplementary Information for "Dynamic land-plant carbon sources in marine sediments inferred from ancient DNA"

### Content


### Supplementary Data

**Supplementary data S1 (as extended data)**

**Supplementary data S2 (as extended data)**

**Supplementary data S3 (as extended data)**

### Supplementary Figures

**Supplementary figure 1**

**Supplementary figure 2**

**Supplementary figure 3**

**Supplementary figure 4**

**Comparison of sedaDNA results of all marine cores to various terrestrial proxies**

**Fram Strait**

**Supplementary Figure 5**

**Off-Kamchatka**

**Supplementary Figure 6**

**Off-Tobago**

**Supplementary Figure 7**

**Supplementary Figure 8**

**Off-Australia**

**Supplementary Figure 9**

**Bransfield Strait**

**Supplementary Figure 10**

**Supplementary figure 11**

### Supplementary references

<u>Supplementary Figures</u>

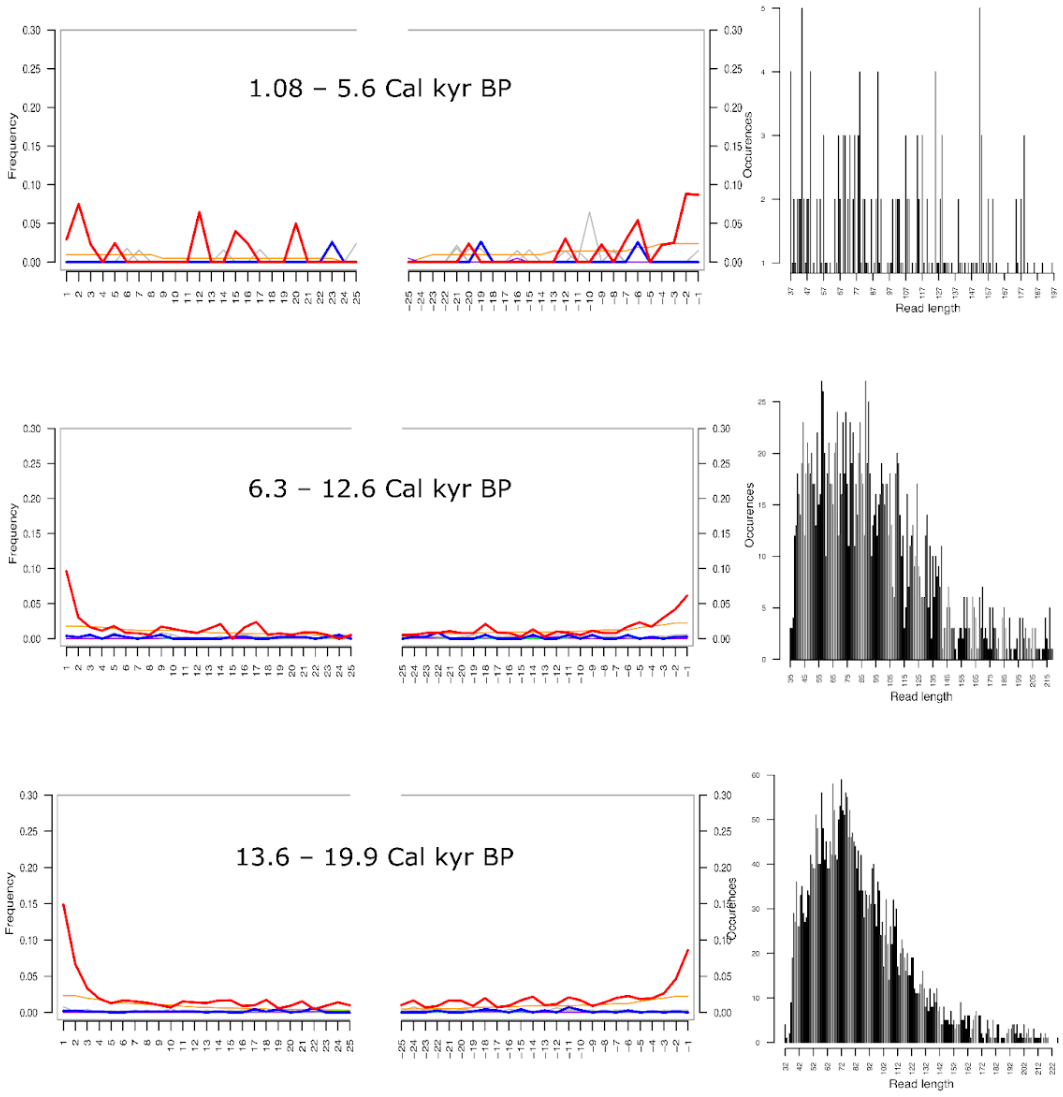


**Supplementary Figure 1: Postmortem DNA damage patterns for *Salix* sp. (TaxID 40685) DNA reads extracted from marine sediments for the core SO201-2-12KL grouped in 3 datasets (ds01: 1081–5600 years BP; ds02: 6200–12,600 years BP; ds03: 13,600–19,900 years BP)**. Reads were mapped against the *Salix brachista* reference genome (Accession number: GCA_009078335.1). Left plots: The red line indicates an increase of misincorporation (C to T changes) due to DNA damage towards the 5ʻ end of the *Salix* DNA reads. The orange line indicates soft clipped bases that did not align to the reference and were excluded from the damage pattern analysis. Right plots: Length distribution of *Salix brachista* reads used for damage pattern analysis.

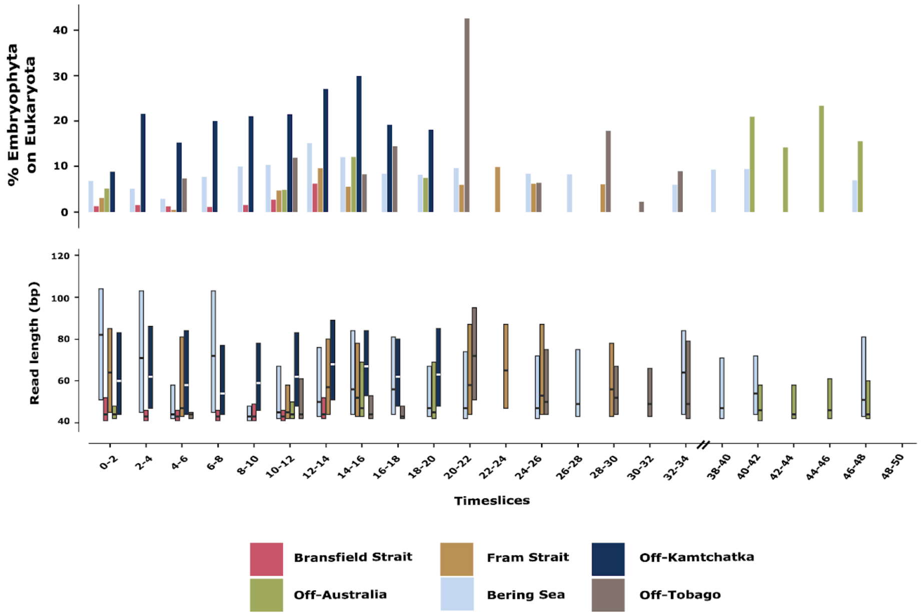


**Supplementary Figure 2: The share of Embryophyta (land plants) on Eukaryota (upper panel) and read length (bp) of Embryophyta sequences (lower panel) of each marine sediment core over 50,000 years in time slices (2000 years).** The boxplots are shown with a coefficient=0 and without outliers. The absence of data in the time slices between 34,000 and 38,000 years ago is indicated by two prominent lines denoting a break in the axis.

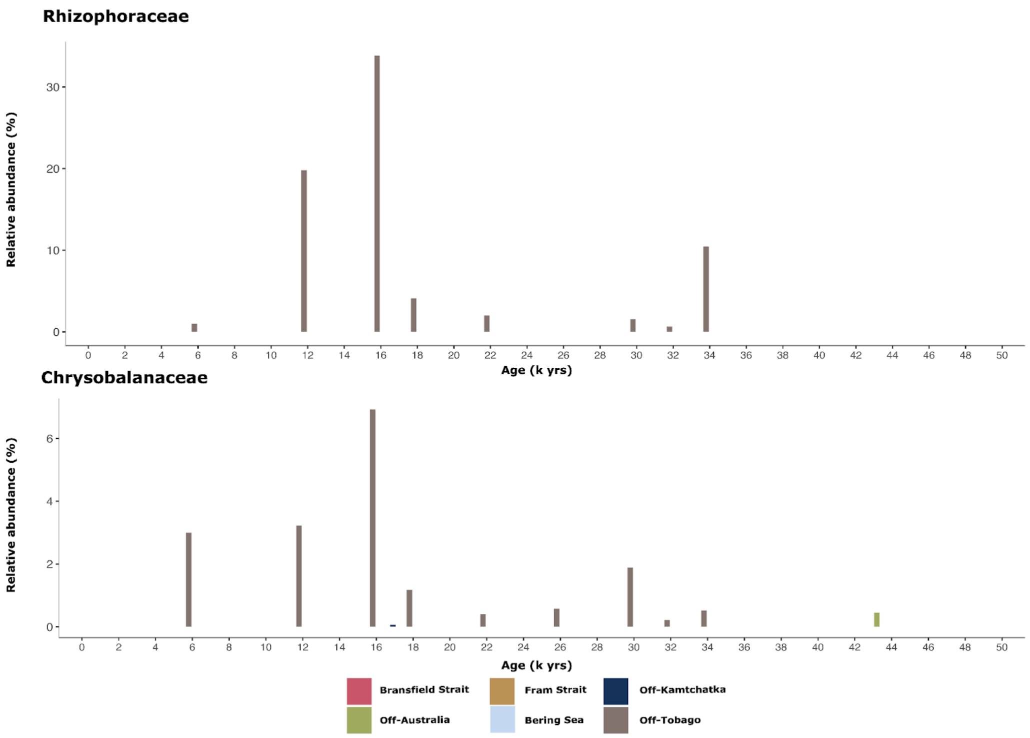


**Supplementary Figure 3: Relative abundance (%) of Rhizophoraceae and Chrysobalanaceae in the sedaDNA dataset from marine sediment cores over time (thousand (k) yrs).**

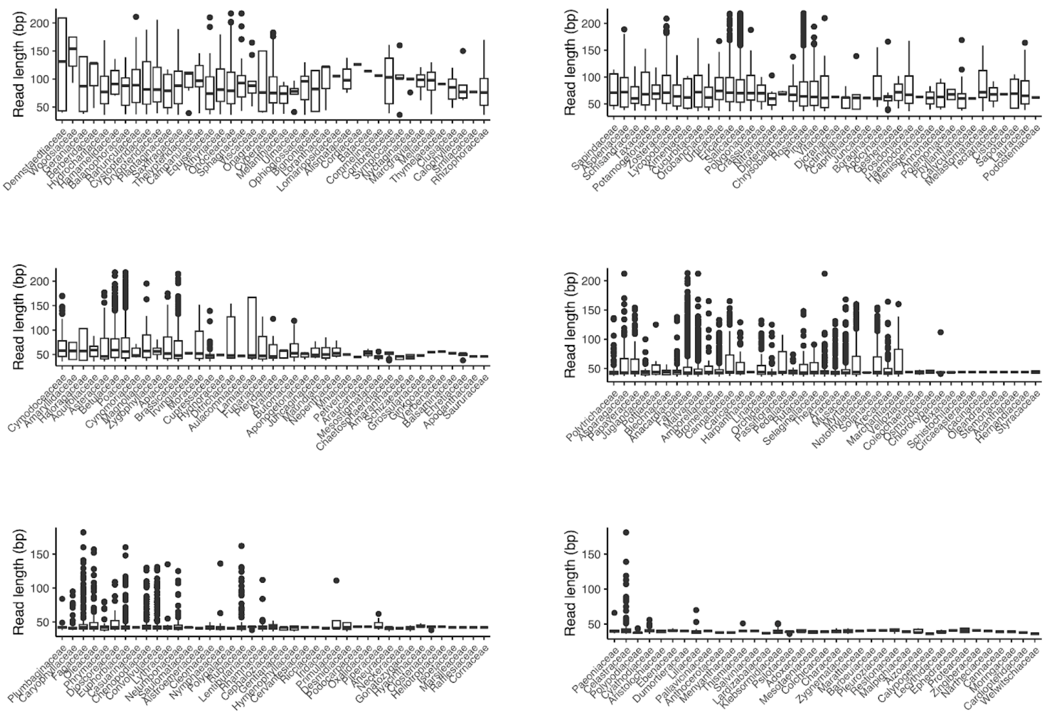


**Supplementary Figure 4: Read length of all 242 Embryophyta families in the sedaDNA dataset for all 6 marine sediment cores together.**

### Comparison of sedaDNA results of all marine cores to various terrestrial proxies

1. Fram Strait

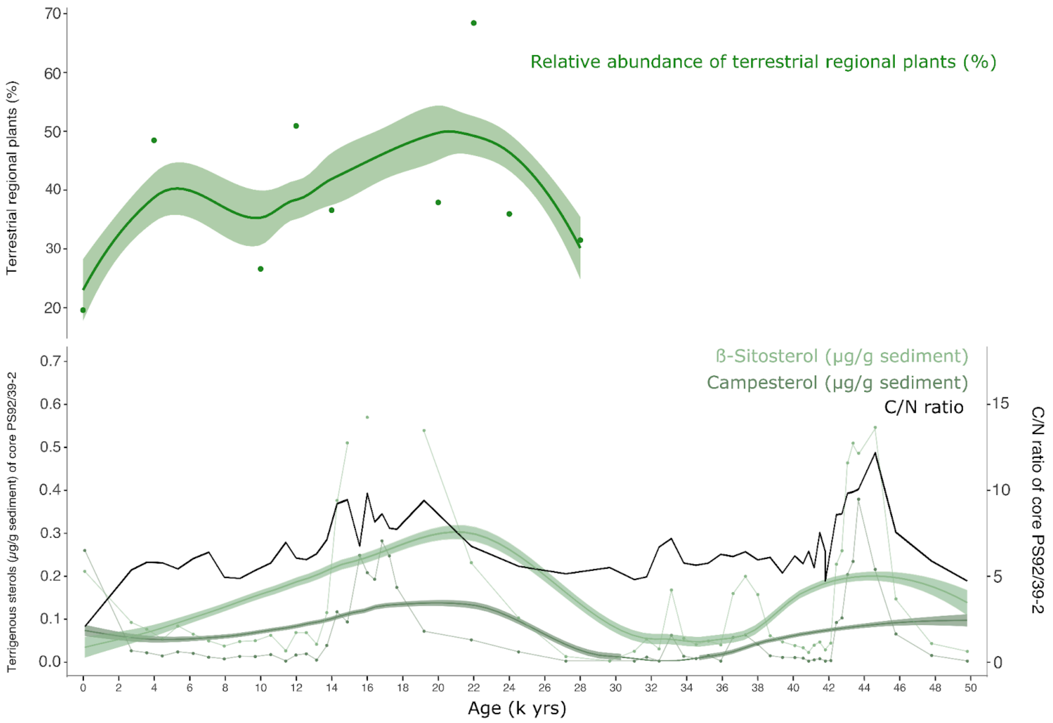


**Supplementary Figure 5: Comparison of the sedaDNA derived relative abundance of terrestrial regional plants (% on Eukaryota) of the Fram Strait with the analysis of terrigenous sterols in a nearby marine sediment core.** The relative abundance of terrestrial regional plants (%) is from the marine sediment core MSM05/5-712-2 in the Fram Strait. The values for ß-sitosterol (µg/g sediment) and campesterol (µg/g sediment) are from a proximal marine sediment core (PS92/039-2), published by Kremer et al., 2018.

The relative abundance of terrestrial regional plants (% on Eukaryota) in our marine sedaDNA dataset (MSM05/5-712-2), the higher land plant sterols ß-Sitosterol and Campesterol, and the C/N ratio all show elevated values during the last deglacial (21–12 k yrs) (Supplementary Figure 5). Core PS92/039-2, from which the land plant sterols and the C/N ratio originate, is located on the Yermak Plateau (Kremer et al., 2018). The values of the terrestrial regional plant DNA fits to the temporal pattern of the C/N ratio and both terrigenous sterols. All proxies show a massive discharge of terrigenous material during the last deglaciation. Additionally the sedaDNA dataset shows the source of this terrestrial organic matter. The source of the terrigenous material is mainly grasslands (Poaceae) and wetland taxa like Salicaceae (Figure 5).

2. **Off-Kamchatka**

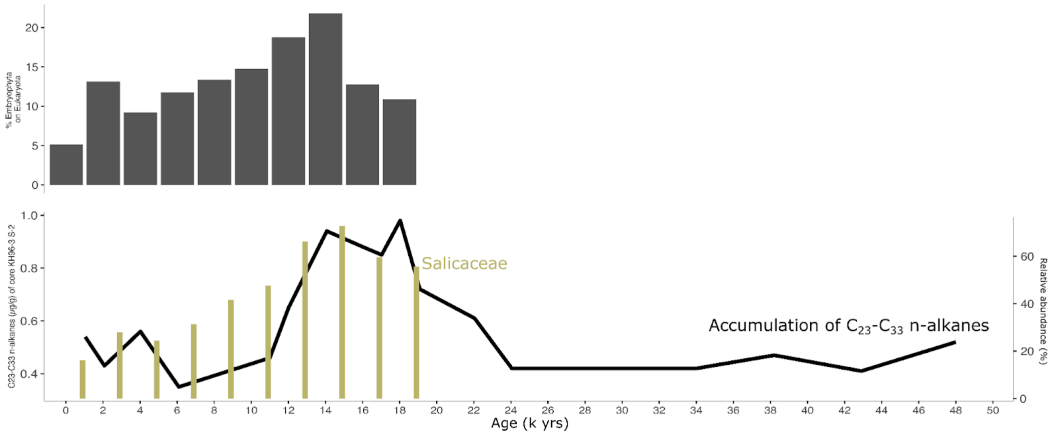


**Supplementary Figure 6: Comparison of Embryophyta on Eukaryota ratio (%), the relative abundance of Salicaceae and the accumulation fo $C_{23}$-$C_{33}$ n-alkanes (Amo & Minagawa, 2003).** The sedaDNA data derived from the marine sediment core SO201-2-12KL from this study. The n-alkanes dataset originates from the proximal KH96-3-S-2 marine sediment core published by Amo & Minagawa, 2003.

Terrestrial biomarkers like n-alkanes can show the accumulation of higher land-plant organic matter (Amo & Minagawa, 2003). As shown in this study there is a stronger terrestrial signal during the deglacial compared to the Holocene (Supplementary Figure 6). Furthermore the sedaDNA results also identify Salicaceae as the main Embryophyta family behind that signal. Members of the Salicaceae family are usually riparian woodland taxa which favor humid weather conditions. We propose that this Salicaceae derived terrestrial organic matter was mainly imported through rivers to the location of the marine sediment core SO201-2-12KL.

3. **Off-Tobago**

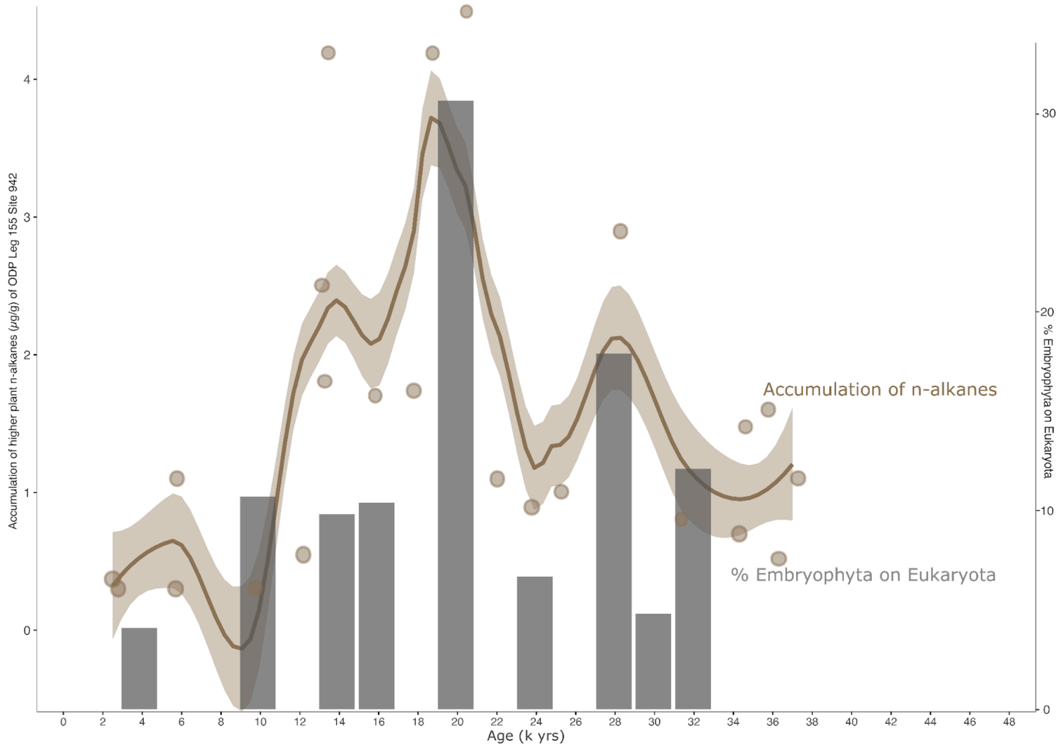


**Supplementary Figure 7: Comparison of Embryophyta on Eukaryota ratio (%) and the accumulation of n-alkanes of higher land plants (Boot et al., 2006).** The sedaDNA data are derived from the marine sediment core of the Tobago Basin M78/1-235-1 from this study. The n-alkanes dataset originates from the proximal ODP Leg 155 Site 942 marine sediment core published by Boot et al., 2006.

The accumulation rate of n-alkanes (µg/g) of the core ODP Leg 155 Site 942 (Boot et al., 2006) and the ratio of Embryophyta on Eukaryota (%) of the core M78/1-235-1 show the same trend (Supplementary Figure 7). The increase in terrestrial material during the Last Glacial and deglacial could be a result of more discharge of the Amazon river but also changes in sea level (Boot et al., 2006).

Furthermore, the comparison of the accumulation of Taraxerol/n-$C_{28}$-ol in core ODP Leg 155 Site 942 and the relative abundance of Rhizophoraceae in core M78/1-235-1 show the same temporal pattern (Supplementary Figure 8). The biomarker Taraxerol/n-$C_{28}$-ol shows the occurrence of mangroves as part of the terrestrial signal (Versteegh et al., 2004). From this, one could conclude that the productivity of the mangroves increased during the deglacial period or that more river runoff occurred due to meltwater and increased precipitation (Boot et al., 2006).

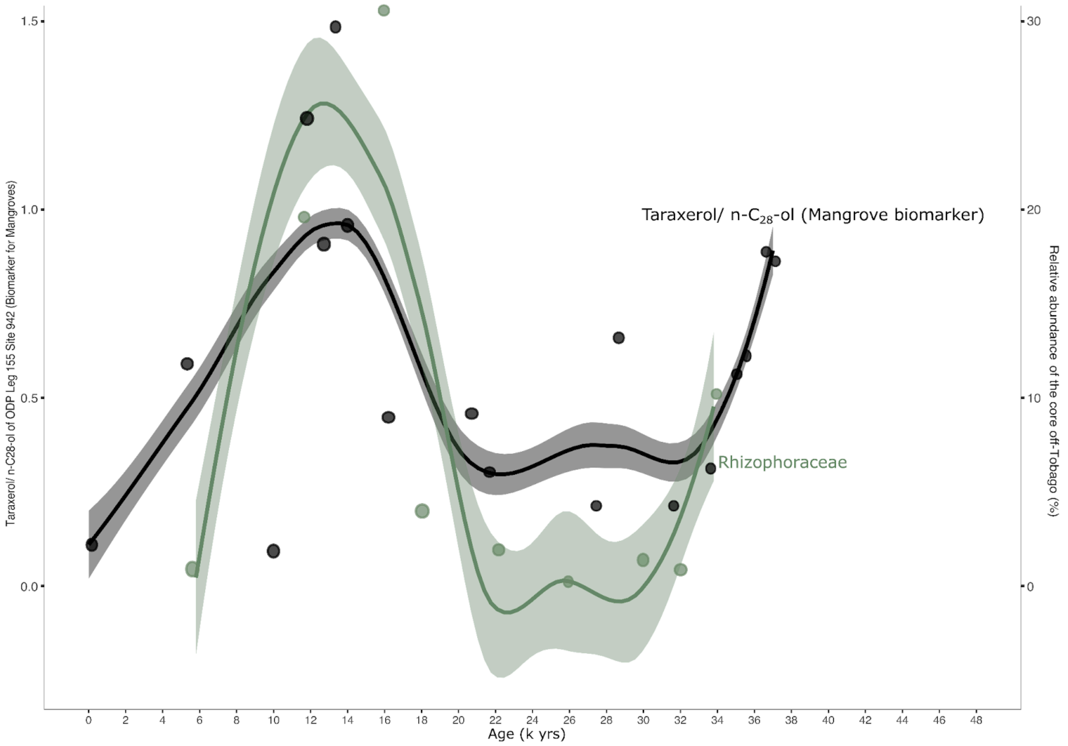


**Supplementary Figure 8: Comparison of relative abundance of Rhizophoraceae (%) and the accumulation of Taraxerol/n-$C_{28}$-ol (mangrove biomarker) (Boot et al., 2006).** The sedaDNA data are derived from the marine sediment core of the Tobago Basin M78/1-235-1 from this study. The Taraxerol/n-$C_{28}$-ol dataset originates from the proximal ODP Leg 155 Site 942 marine sediment core published by Boot et al., 2006.

4. **Off-Australia**

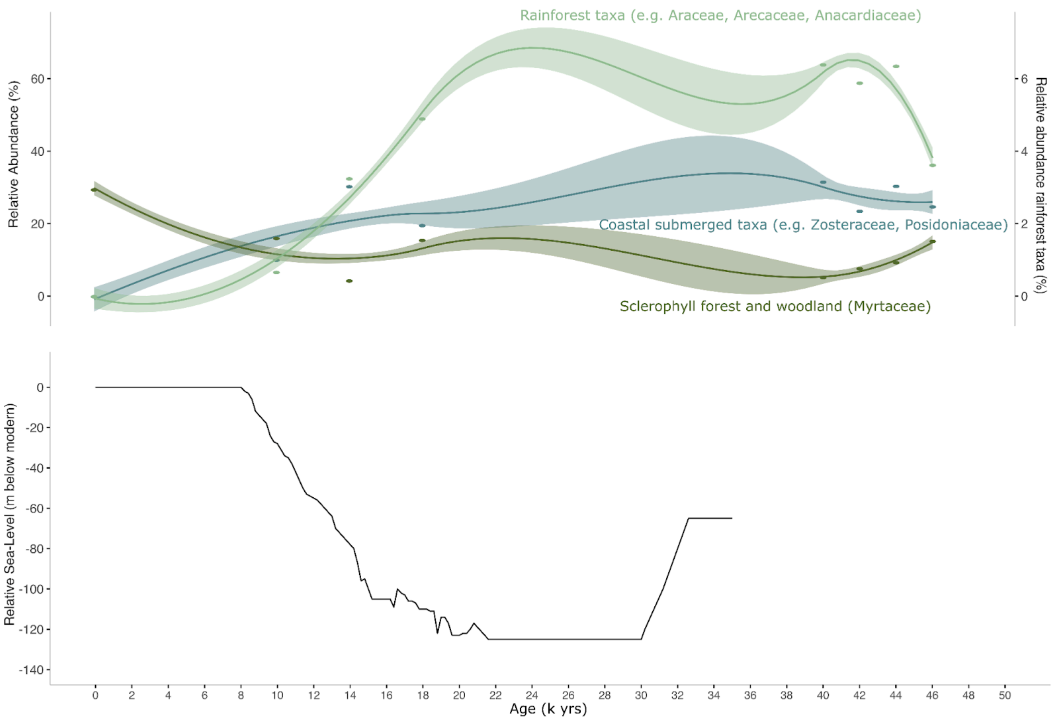


**Supplementary Figure 9: Relative contribution (%) of coastal submerged taxa and terrestrial regional taxa in the Great Australian Bight compared to relative sea level (m below modern).** The sedaDNA data are derived from the marine sediment core MD03-2614G located on the south eastern coast of Australia in the Great Australian Bight. The terrestrial regional taxa are separated into Sclerophyll forest and woodland vegetation (Myrtaceae) and Rainforest taxa (Araceae, Arecaceae, Anacardiaceae). The data on the relative sea level (m below modern) were published by Lewis et al., in 2013.

The sedaDNA dataset of the marine sediment core MD03-2614G shows a clear shift between the contribution of coastal submersed plants such as Zosteraceae, Cymomodaceae, and Posidoniaceae and terrestrial regional taxa such as Myrtaceae to the organic matter of the marine sediment core. Since 34,000 cal yr BP, the relative abundance of coastal submerged taxa decreased and terrestrial regional taxa became more abundant (Supplementary Figure 9). We hypothesize a connection with the changes in sea level since the last glacial and the increased meltwater runoff during the deglacial, which increased river discharge (Lewis et al., 2013). In this case, the sedaDNA dataset allows us to directly compare the contribution of coastal submerged and terrestrial land plants to sedimented organic matter. In addition, our shotgun data show a decline in rainforest taxa from the last glacial period to the Holocene. The pollen-derived precipitation pattern from Dekker et al. (2019) shows the same trend on the southwest coast of Australia.

5. **Bransfield Strait**

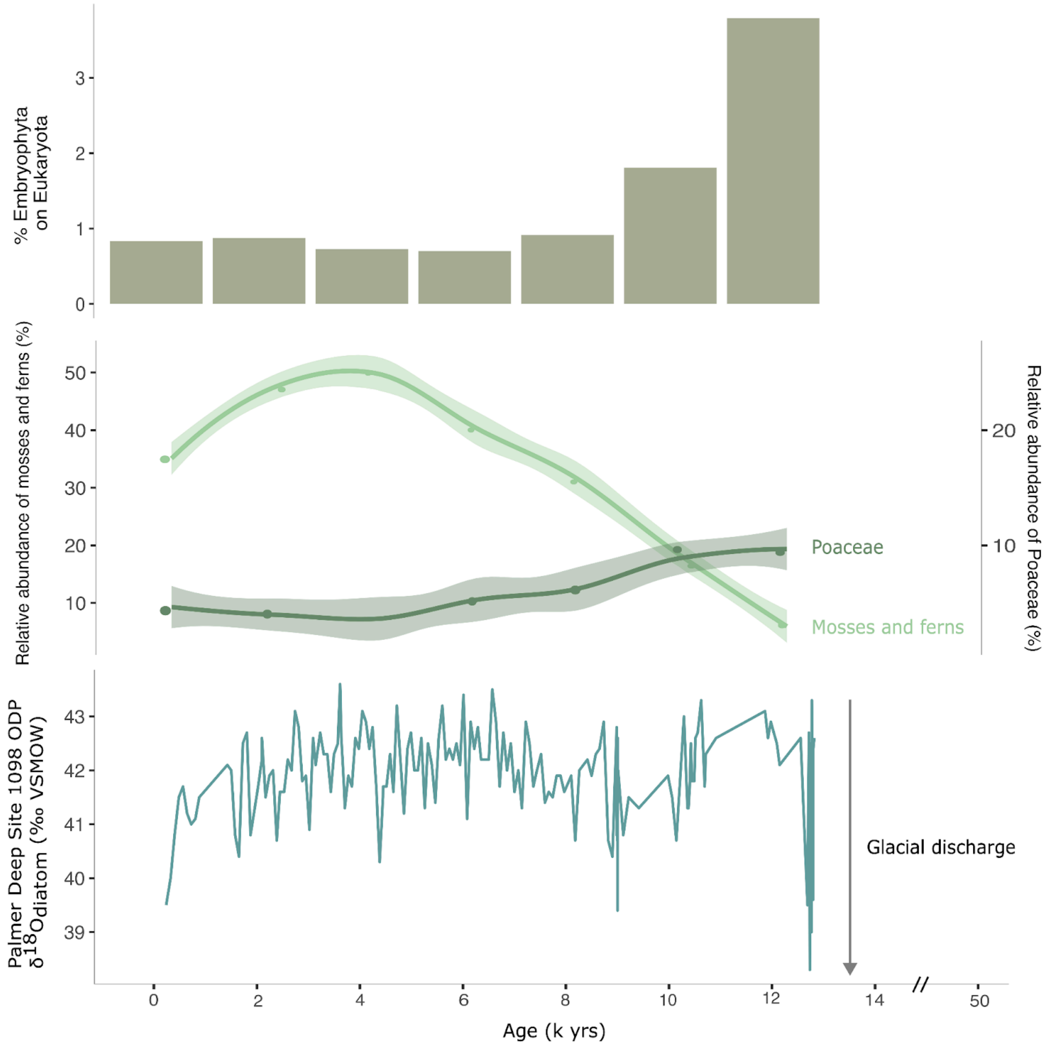


**Supplementary Figure 10: Comparison of Embryophyta on Eukaryota ratio (%), the relative abundance of mosses and ferns and Poaceae (%) with the $\delta^{18}O_{diatom}$ (‰ VSMOW) values of the marine sediment core ODP site 1098 published by Pike et al., 2013.** The x-axis has a break between 14 cal kyr BP and 50 cal yr BP. The $\delta^{18}O_{diatom}$ (‰ VSMOW) record reveals the amount of glacial discharge.

The terrestrial input to the Southern Ocean is low (1–3%) compared to the tropics (5–40%) and the Northern Hemisphere (5–23%) (Fig. 2). However, the SedaDNA dataset shows a temporal pattern of Embryophyta DNA in the Bransfield Strait record (Supplementary Figure 10). More terrestrial material is introduced into the Bransfield Strait during the last deglacial. Between 14 cal kyr BP and 12 cal kyr BP, glacial discharge increased significantly near the Bransfield Strait in the Palmer Deep (Pike et al., 2013) (Supplementary Figure 10). We hypothesize that this is the same reason for the higher input of terrestrial land plant DNA in the Bransfield Strait.

Based on the Embryophyta DNA dataset, we see that most of the terrestrial DNA consists of short and highly degraded fragments that are erroneously assigned to the overrepresented crop plants (Solanaceae, Fabaceae) (Fig. 2). Nevertheless, there is also a high proportion of Poaceae at this time, which fits the dry and cold conditions (Ingolfsson et al., 2003) (Supplementary Figure 10).

It is known that moss beds formed on the Antarctic Peninsula and the surrounding islands after the retreat of the glaciers in the Holocene due to the increasing warmth and humidity (Ingolfsson et al., 2001). The sedaDNA dataset from the marine sediment core PS97/72-01 also demonstrates a strong increase in mosses throughout the Holocene. However, due to the lower glacial discharge, less terrestrial plant material was deposited in the marine sediments in total (Supplementary Figure 10).

POST HOC TESTS

Games-Howell Post-Hoc Test – Read_Length

| | | Bering Sea | Bransfield Strait | Fram Strait | off-Australia | off-Kamchatka | off-Tobago |
|---|---|---|---|---|---|---|---|
| Bering Sea | Mean difference | – | 17.85275 | -0.2296210 | 11.241775 | -8.480219 | -3.525913 |
| | p-value | – | < .0000001 | 0.0010311 | < .0000001 | < .0000001 | < .0000001 |
| Bransfield Strait | Mean difference | | – | -18.0823696 | -6.610973 | -26.332968 | -21.378662 |
| | p-value | | – | < .0000001 | < .0000001 | < .0000001 | < .0000001 |
| Fram Strait | Mean difference | | | – | 11.471396 | -8.250598 | -3.296292 |
| | p-value | | | – | < .0000001 | < .0000001 | < .0000001 |
| off-Australia | Mean difference | | | | – | -19.721994 | -14.767688 |
| | p-value | | | | – | < .0000001 | < .0000001 |
| off-Kamchatka | Mean difference | | | | | – | 4.954306 |
| | p-value | | | | | – | < .0000001 |
| off-Tobago | Mean difference | | | | | | – |
| | p-value | | | | | | – |

**Supplementary Figure 11: Results from Welch's test and Games-Howell post-hoc test evaluating differences in read length between the marine sediment cores.** We performed Welch's test to accommodate unequal variances and subsequently employed the Games-Howell post-hoc test to identify significant group differences.
